# Long noncoding RNAs sustain high expression of exogenous Oct4 by sponging miRNA during reprogramming

**DOI:** 10.1101/612077

**Authors:** Qingran Kong, Xiaolei Zhang, Jiaming Zhang, Kailun Zheng, Heng Zhang, Xixiang Pei, Zhi Yin, Duancheng Wen, Zhonghua Liu

**Affiliations:** Key Laboratory of Animal Cellular and Genetics Engineering of Heilongjiang Province, College of Life Science, Northeast Agricultural University, Harbin, 150030, Heilongjiang Province, China; Ronald O. Perelman and Claudia Cohen Center for Reproductive Medicine, Weill Cornell Medical College, New York, NY 10065, USA; Harbin Botai Bio-Tech Co., Ltd., No. 59 mucai Street, Xiangfang District, Harbin, 150030, Heilongjiang Province, China

**Keywords:** ceRNAs, circRNAs, lincRNAs, miRNAs, reprogramming

## Abstract

Long noncoding RNAs (lncRNAs) modulate gene expression as competing endogenous RNAs (ceRNAs) via sponging microRNAs (miRNAs). However, the extent and functional consequences of ceRNAs in diverse cellular context still need to be proven. Using a doxycycline inducible expression of Yamanaka four factors to generate induced pluripotent stem cells (iPSCs) from mouse embryonic fibroblasts (MEFs), we found the miRNAs from MEFs remained highly expressed from day 0 to day 6 after doxycycline induction; unexpectedly, many genes targeted by these miRNAs were actually up-regulated; meanwhile, long intergenic noncoding RNAs (lincRNAs) and circular RNAs (circRNAs) which have complementary binding sites with the miRNAs were highly expressed, indicating lincRNAs and circRNAs (linc/circRNAs) may serve as sponges for miRNAs to block their activities during reprogramming. Intriguingly, the knockdown of the linc/circRNAs sponging the miRNAs targeting Oct4 mRNA resulted in down-regulation of exogenous Oct4 expression, decrease of reprogramming efficiency, and low-grade chimera forming iPSCs. Our results suggest that the ceRNA network plays an important role in reprogramming somatic cells to pluripotent stem cells.

## Introduction

The microRNAs (miRNAs, ~22nt noncoding RNAs) are an abundant class of small noncoding RNAs, which guide the RNA-induced silencing complex (RISC) to miRNA response elements (MREs) to its targeting transcripts, and can induce mRNA degradation or translational repression [1–3]. The target efficiency of miRNAs is known related to the abundance of MREs present in the transcripts [4–6]. Previous studies have shown that one gene can have multiple MREs for distinct miRNAs, and vice versa, miRNA can target multiple distinct transcripts [7, 8]. A highly expressed MREs-containing transcript can compete for multiple shared miRNAs and lead to observable changes in miRNA activity and thus regulate the expression of one or multiple miRNA target transcripts. This “sponge” mechanism has been proposed for competing endogenous RNAs (ceRNAs) to regulate other RNA transcripts by competing for shared miRNAs [9, 10].

Such sponge mechanism involves coding and noncoding transcripts, including long intergenic noncoding RNAs (lincRNAs) [10–13]. For example, linc-MD1, a muscle-specific ceRNA, regulates muscle differentiation [13]. Recently, another naturally occurring family of noncoding RNAs, the circular RNA (circRNA), from back-spliced exons, has also been reported to act as miRNA sponges to regulate gene expression [14–17], such as miR-7 inhibitor in the central nervous system (CNS) [15]. Though thousands of lincRNAs have been annotated in eukaryotic genomes and targeted by miRNAs with both computational and experimental evidences [18, 19], only a small number of lincRNAs have been identified to serve as ceRNAs so far. The circRNAs are also found abundant in the eukaryotic transcriptome [20, 21], whether circRNAs can function as miRNA sponge remains completely elusive. Moreover, the biological relevance of ceRNAs has recently been challenged, because the relatively low expression level of most lncRNAs might limit their ability to effectively modulate, in a miRNA-dependent manner, mRNA abundance. For example, the level of one transcript, Aldoa, required to significantly alter the level of one highly abundant miRNA, miR-122, and its targets in adult hepatocytes was found to exceed the changes observed in vivo, even under extreme physiological or disease conditions [5]. The discrepancies in the conclusions of the different attempts suggest that substantially more genetic and genomic evidence from diverse cell types will be required to resolve this issue and to establish the general prevalence and physiological relevance of ceRNAs.

Considering that many established ceRNAs neither share an unusual high number of predicted MREs with their mRNA targets nor are especially abundant [10, 22], the ability to modulate mRNA abundance was thought to be limited [4, 6]. However, accumulating evidences suggest that ceRNAs might play an important role in regulating key transcription factors during cell-fate decision process, supported by the observation that the expression of ceRNAs is tightly regulated with spatial and temporal expression patterns. For example, linc-RoR competes for miR-145 binding with key self-renewal transcription factor transcripts, including Nanog, Oct4, and Sox2, and is expressed in undifferentiated embryonic stem cells (ESCs) [11]. The specific expression profile of these ceRNAs might specify the cells in which their activity exerts the greatest effect. Cell-fate decisions often involve switch-like responses in the expression levels of key regulatory genes that result in coordinated changes in transcription profiles, suggesting that relatively lowly abundant, yet specifically expressed, ceRNAs might function efficiently on regulation of the transition from differentiated to pluripotent cell states.

Cell fate conversion can be induced by over-expressing sets of regulatory transcription factors [23–25]. The over-expression of Oct4, Sox2, Klf4 and c-Myc (OSKM) can reprogram mouse embryonic fibroblasts (MEFs) to induced pluripotent stem cells (iPSCs) [26, 27]. In the study, we performed ribo-minus RNA-seq and small RNA-seq from cells collected at the early reprogramming stages and fully reprogrammed iPSCs which can generate “all-iPSCs” pups (Fig 1A), to profile expression patterns of miRNAs, lincRNAs, circRNAs and mRNAs. We also took advantage of publicly available RNA-seq and small RNA-seq data from Mbd3^flox/-^ reprogramming system, which results in near 100% efficiency of iPSCs reprogramming within 7 days by OSKM induction [28], to confirm our analysis. Our studies ought to determine the possible function of linc/circRNAs as miRNAs sponge during cellular reprogramming.

**Figure 1.**
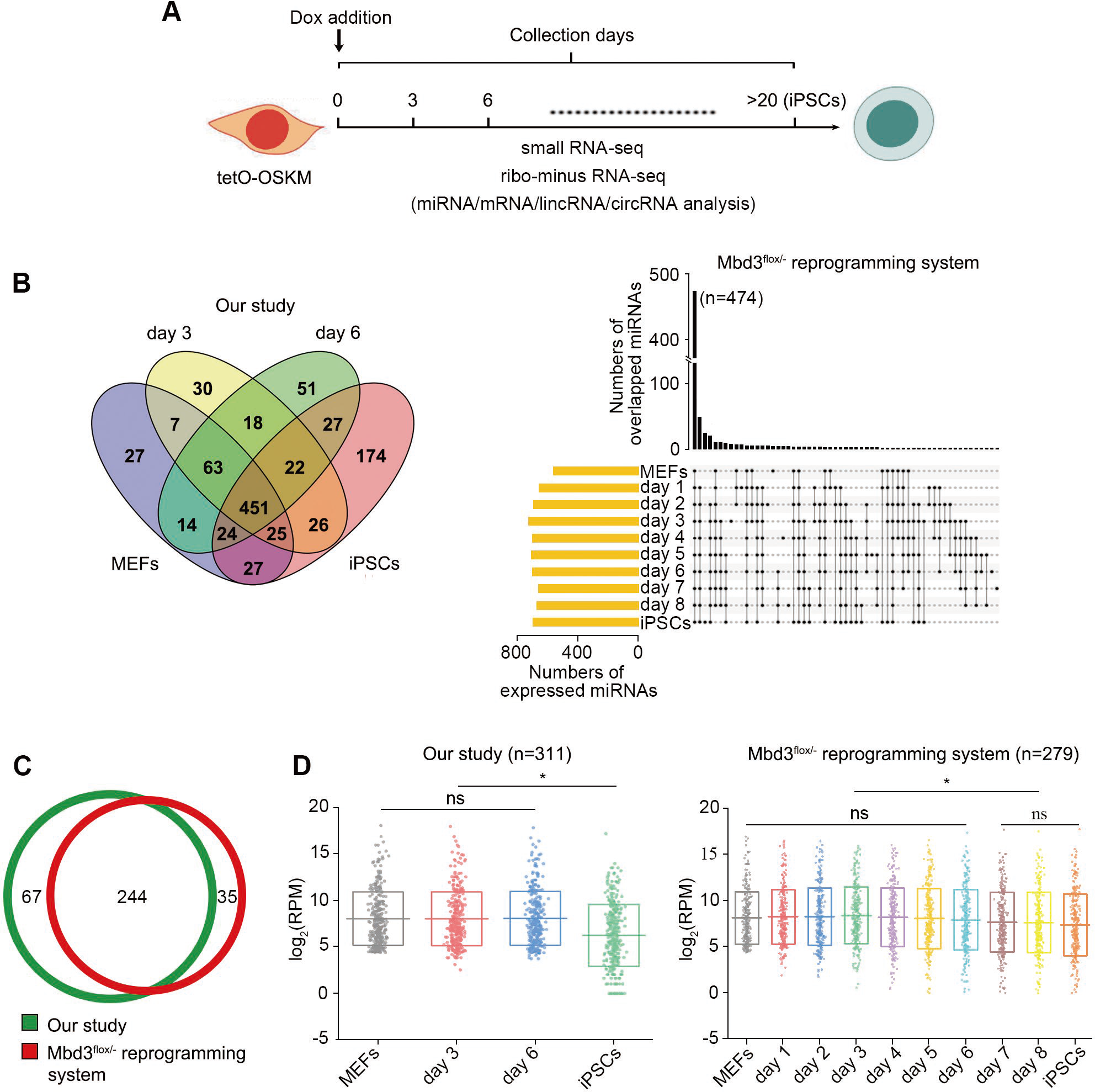
Profiling of miRNAs abundance during reprogramming. A Schematics of iPSCs reprogramming indicating the time-points at which samples were collected for libraries preparation. B Overlap of miRNAs expressed in different reprogramming stages and fully reprogrammed iPSCs. C Highly-expressed miRNAs in MEFs. Only the top 50% abundant and RPM > 20 miRNAs are selected. 244 miRNAs were overlapped between the two reprogramming systems. D Expression patterns of the MEFs-highly expressed miRNAs over the course of reprogramming and in iPSCs. * indicates p <0.05.

## Results

### Profile of miRNAs from MEFs during the course of reprogramming

MEFs from mice carrying a single dox-inducible OSKM poly cistronic 4F2A cassette (tetO-OSKM) were induced reprogramming by adding doxycycline to the culture medium. To demonstrate the miRNA-mediated crosstalk during reprogramming, we first profiled small RNAs in MEFs, the reprogramming MEFs (rMEFs) at day 3 and day 6 post-OSKM induction and the fully reprogrammed iPSCs. To confirm our data, profiling of small RNA from Mbd3^flox/-^ reprogramming system were also analyzed. 451 and 474 miRNAs were co-expressed among MEFs, rMEFs and iPSCs in our and Mbd3^flox/-^ reprogramming systems, respectively, indicating miRNAs from MEFs are sustainable during reprogramming (Fig 1B). To figure out the extent of miRNA-mediated interactions between their mRNA targets and lncRNAs, we selected311 and 279 miRNAs highly expressed in our and Mbd3^flox/-^MEFs (top 50% abundant and RPM >20), and 244 miRNAs were overlapped (Fig 1C). The expression of the MEFs-highly expressed miRNAs were not significantly changed over the course of reprogramming, and significantly decreased in reprogrammed cells (at day 7 and day 8 in Mbd3^flox/-^ reprogramming system) and iPSCs (Fig 1D; Table EV1), and the miRNAs were chose for further analysis.

### Activity of miRNAs may be inhibited during reprogramming

During reprogramming, genes associated with pluripotent establishment need to be activated or up-regulated, whereas developmental genes need to be silenced [28]. In order to determine the role of miRNAs in regulating the transcriptomic changes, we profiled expression of mRNAs on MEFs, rMEFs, and iPSCs as well, and then analyzed the effect of the MEFs-highly expressed miRNAs on up-and down-regulated genes during reprogramming. We found the miRNAs not only targeted the down-regulated genes, but also the up-regulated genes, including key pluripotent transcriptional factors, such as Oct4, Klf4, Nanog, Tbx3, Sall4 and Esrrb. Notably, the number of up-regulated genes targeted by the miRNAs was more than that of the down-regulated ones (Fig 2A and Table EV2). And, the expression levels of miRNAs targeting up- and down-regulated genes had no significant difference (*p* >0.05), and both maintained at high levels during the course of reprogramming and decreased in iPSCs (Fig 2B). Then, we checked the densities of MREs in up-and down-regulated mRNA targets. We found that the MRE density in down-regulated mRNA targets (mean of 0.430 MREs/kb) is not significantly different with up-regulated mRNA targets (mean of 0.487 MREs/kb; *p*>0.05) (Fig 2C). Considering that high-affinity MREs would be more effective to bind miRNAs than low-affinity MREs, resulting in down-regulation of gene expression more efficiently, we analyzed the ratio of 8mer, 7mer and 6mer MREs in the targets, and found no significant difference between the up-and down-regulated targets (Fig 2D). Our results show that neither the density nor the affinity of MREs is different between the miRNAs targeted up-and down-regulated genes. During reprogramming, developmental genes should be silenced, so the highly-expressed miRNAs targeting the down-regulated genes may have little effect on reprogramming. However, if the expression of the up-regulated genes is influenced by the miRNAs, the reprogramming will fail. Thus, we hypothesize that additional mechanisms may exist to inhibit the activity of the MEFs-highly expressed miRNAs during reprogramming.

**Figure 2.**
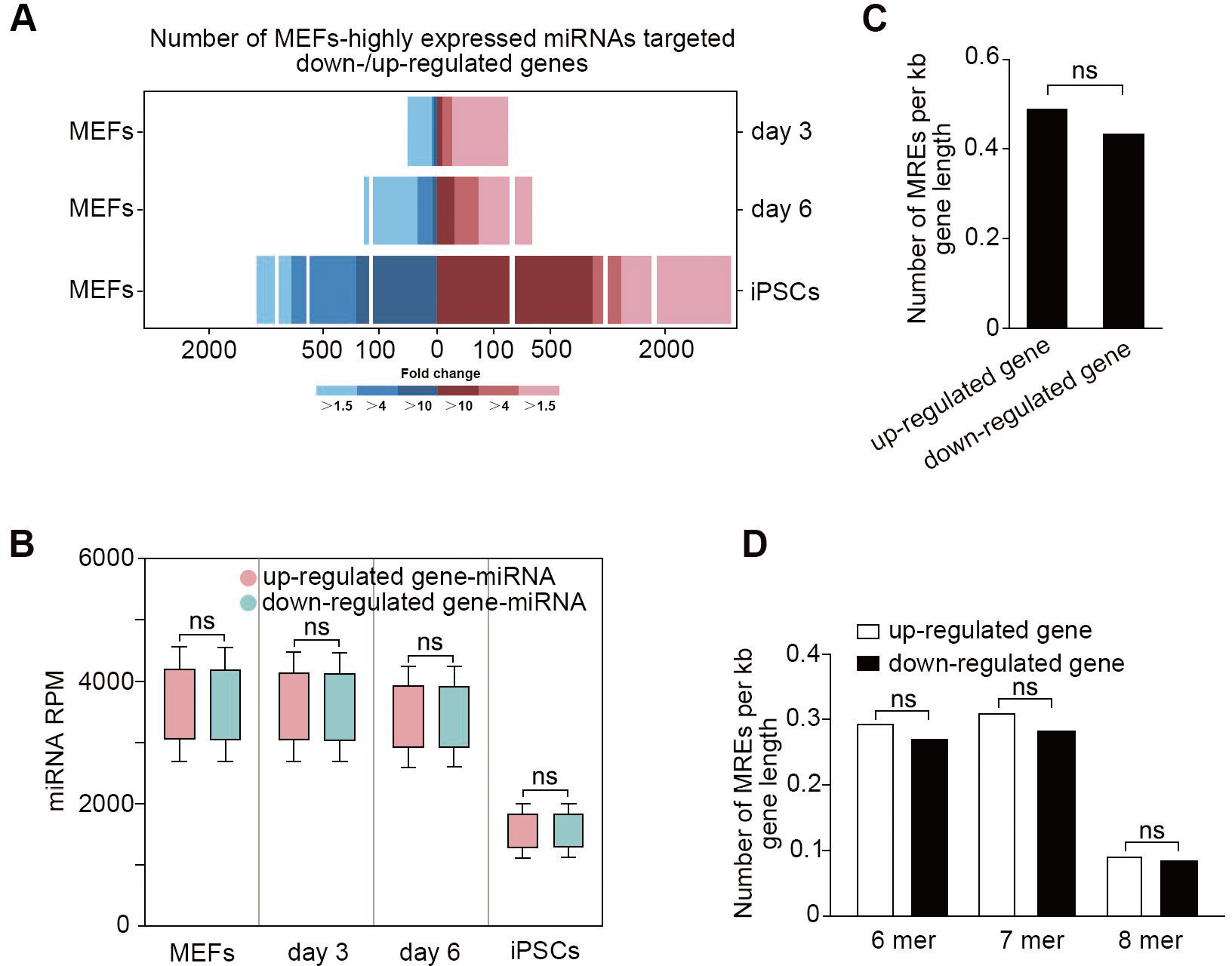
Activity of miRNAs may be reduced during reprogramming. A The numbers of differentially expressed genes targeted by MEFs-highly expressed miRNAs between MEFs and rMEFs or iPSCs. Blue and red bars represent down- and up-regulated genes targeted by the miRNAs, respectively. B Expression patterns of miRNAs targeting down- and up-regulated genes during reprogramming. There is no significant difference in expression between the two kinds of miRNAs. Up-regulated gene-miRNA: miRNA targeting on up-regulated gene; down-regulated gene-miRNA: miRNA targeting on down-regulated gene. The error bars represent s.d. C Densities of MREs in up-regulated and down-regulated mRNA targets. There is no significant difference for MREs densities between down-regulated (mean of 0.430 MREs/kb) and up-regulated (mean of 0.487 MREs/kb; *p* >0.05). D Analysis of MREs affinity in up-regulated and down-regulated mRNA targets. The ratio of 8 mer, 7 mer and 6 mer MREs in up-regulated targets is 0.092, 0.312 and 0.292 MREs/kb, and 0.084, 0.282 and 0.269 MREs/kb in down-regulated targets, respectively.

### Linc/circRNAs may serve as sponges for multiple miRNAs during reprogramming

Previous studies have shown that linc/circRNAs can function as miRNA sponges to reduce miRNA activity to regulate its targets [11, 14]. We propose that linc/circRNAs may serve as sponges to counteract the activities of the MEFs-highly expressed miRNAs during reprogramming. Taking advantage of ribo-minus RNA-seq, we characterized linc/circRNAs in rMEFs and iPSCs. 1490 known lincRNAs and 641 known circRNAs were identified (Table EV3). Comparing to MEFs, 101 lincRNAs (Fig 3A) and 204 circRNAs (Fig 3B) were found to express with higher levels in day 3 and day 6 rMEFs. The limitation of sequencing libraries from Mbd3^flox/-^ reprogramming system is the inability to detect non polyadenylated (polyA-) RNA, so we cannot identify circRNAs using the data. However, 93 highly-expressed lincRNAs were characterized during the course of the reprogramming (Fig 3A). Strikingly, we found that almost all of the MEFs-highly expressed miRNAs shared complementary binding sites with the highly-expressed linc/circRNAs with only few exceptions (Fig 3C), indicating that linc/circRNAs might regulate gene expression through inhibiting the activity of miRNAs during reprogramming. In addition, we observed that some miRNAs, proven to serve as reprogramming barriers, such as let-7 and miR-34 families [29, 30], were highly expressed during reprogramming, and their expressions were decreased in iPSCs (Fig 3D). We wonder if there are linc/circRNAs which can counteract the activity of the miRNAs. As expected, some linc/circRNAs which have complementary binding sites with the miRNAs were highly expressed during reprogramming (Fig 3E). The results suggest that linc/circRNAs may serve as sponges for miRNAs to block their activities during reprogramming.

**Figure 3.**
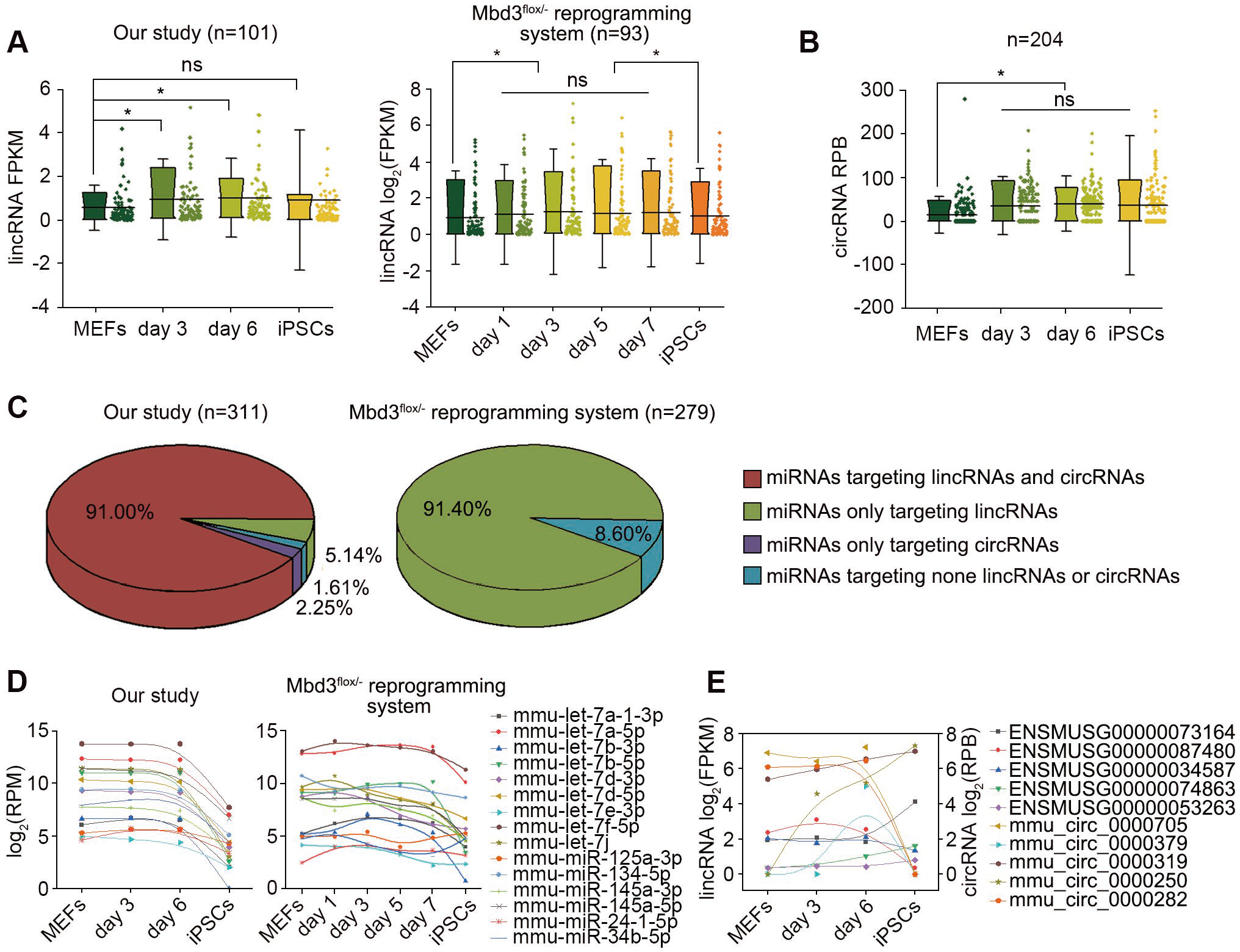
Linc/circRNAs predicted to sponge multiple miRNAs during reprogramming. A LincRNAs highly expressed during the course of iPSCs reprogramming. B CircRNAs highly expressed during the course of iPSCs reprogramming.* indicates p <0.05. The error bars represent s.d. C MEFs-highly expressed miRNAs predict to targeting linc/circRNAs. Almost all of the miRNAs shared complementary binding sites with the highly expressed linc/circRNAs with only few exceptions. D Expression patterns of miRNAs that have been proven to inhibit cellular reprogramming. E Expression patterns of linc/circRNAs predicted to be targeted by the miRNAs inhibiting cellular reprogramming.

### Activity of miRNAs targeting Oct4 is counteracted during iPSCs reprogramming

To further demonstrate the ceRNA mechanism during reprogramming, we tested the hypothesis in the context of Oct4, a key pluripotent transcriptional factor that determines the successful reprogramming [31]. We predicted seven MEFs-highly expressed miRNAs targeting Oct4 mRNA (Oct4-miRNAs) by RNA22 [32] (Fig 4A). However, definitely, Oct4 was up-regulated (Fig 4B). During iPSCs reprogramming, endogenous Oct4 activation generally takes place at 7-12 days [33], and, in the study, we found the expression of endogenous Oct4 was initiated from day 7 post-OSKM induction (Fig 4C), so the ceRNA mechanism we checked mainly involving in the regulation of exogenous Oct4 expression. During the course of reprogramming, the expression of the seven Oct4-miRNAs sustained in high levels, and significantly decreased in reprogrammed cells (at day 7 and day 8 in Mbd3^flox/-^ reprogramming system) and iPSCs (Fig 4D). This result suggests that the expression of Oct4 is not tightly regulated by the miRNAs or a mechanism that can counteract the function of the miRNAs. To confirm if Oct4-miRNAs can down-regulate the expression of exogenousOct4 during the process of reprogramming, the increasing concentrations of the mimics of the miRNAs were transfected into MEFs and the expression of Oct4 mRNA was monitored by qRT-PCR at day 3 post-OSKM induction. We found thatmmu-miR-21a-3p, mmu-miR-421-3p, mmu-miR-497a-5p and mmu-miR-532-3p could efficiently down-regulate Oct4 expression at high concentrations, and the dose-dependent manner revealed that the activity of the miRNAs might be limited (Fig 4E). To ensure that the miRNAs can definitely function on Oct4 mRNA, we inserted the native and mutant MREs of the four Oct4-miRNAs into 3’UTR of a standard luciferase reporter (Fig 4F) and transfected into MEFs, respectively. The luciferase assay was detected at day 3 post-OSKM induction and showed that the four miRNAs indeed functioned on the native MREs but not the mutant ones (Fig 4G). These results suggest that the Oct4-miRNAs can down-regulate exogenousOct4 expression, however, their activities are counteracted during reprogramming.

**Figure 4.**
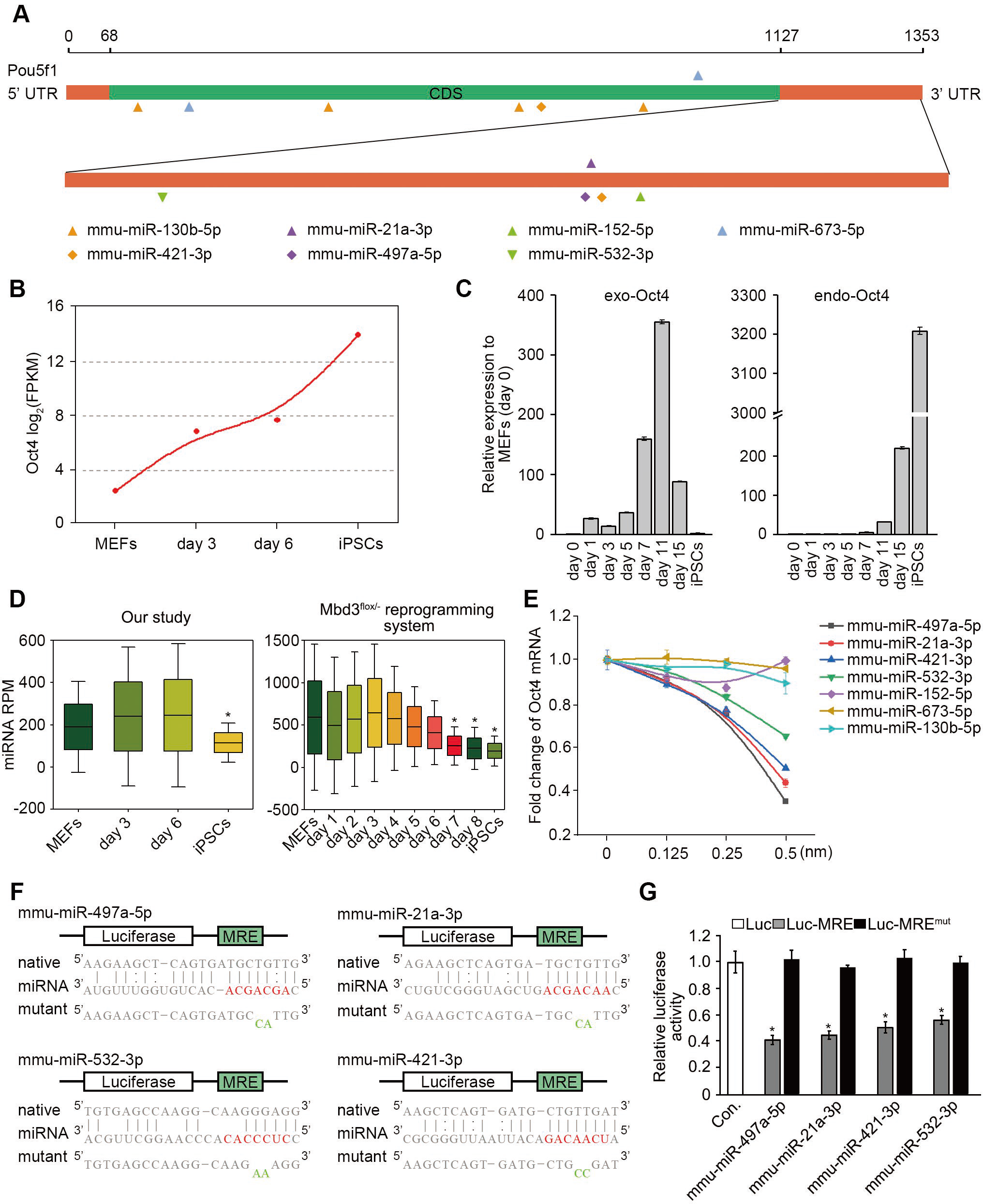
Activity of Oct4-miRNAs is limited during iPSCs reprogramming. A MEFs-highly expressed miRNAs predicted to target Oct4 mRNA (Oct4-miRNAs). Each shape represents a miRNA. B Expression patterns of Oct4 during reprogramming. The expression of Oct4 is shown by red dots. C Expression of exogenousOct4 (exo-Oct4) and endogenous Oct4 (endo-Oct4) during reprogramming checked by qRT-PCR. D Expression patterns of Oct4-miRNAsduring reprogramming.* indicates p <0.05. E Fold change of Oct4 mRNA with increasing concentrations of Oct4-miRNAs as indicated. The increasing concentrations of the miRNAs mimics were transfected into MEFs and expression level of Oct4 mRNA was analyzed by qRT-PCR at day 3 post-OSKM induction. F Luciferase reporter constructs containing the native and mutant MREs in the luciferase 3’UTR. The seed sequences of the miRNAs are highlighted in red, and the mutant nucleotides are shown in green. G Luciferase assay results detected at day 3 post-OSKM. The results showed the four miRNAs functioned on the native MREs but not the mutant ones.* indicates p <0.01. The error bars represent s.e.m.

### Linc/circRNAs can sustain high expression of exogenous Oct4 by sponging the miRNAs targeting Oct4

We considered if the linc/circRNAs which have complementary binding sites with the Oct4-miRNAs could counteract the activity during reprogramming. We indeed found that some linc/circRNAs which have complementary binding sites with the four Oct4-miRNAs were highly expressed during the course of reprogramming, and decreased in reprogrammed cells and iPSCs (Fig 5A). We postulate that if we knockdown the expression of the linc/circRNAs, exogenousOct4 could be down-regulated for the increased activity of the miRNAs. We selected the efficient Locked Nucleic Acid (LNA)-siRNA for each lincRNA and circRNA (LNA-siRNAs targeting the back-splice sequences of circRNAs were used; Fig EV1). As expected, the Oct4 expression was significantly down-regulated at least 20% when some linc/circRNAs were knocked down (Fig 5B). We supposed that these linc/circRNAs might benefit Oct4 expression through sponging the Oct4-miRNAs. To test the hypothesis, we conducted RNA immunopreciptation (RIP) using day3 rMEFs from MEFs transfected with the biotin-labeled miRNAs at a final concentration of 20nM, and observed that ENSMUSG00000092341, mmu_circ_00006895and mmu_circ_00000319were specifically enriched by qRT-PCR analysis normalized to captured Oct4 mRNA (up to 20-fold enrichment compared to gapdh mRNA), suggesting the three linc/circRNAs are able to interact with the Oct4-miRNAs (Fig 5C). Further analysis showed multiple complementary binding sites withmmu-miR-21a-3p, mmu-miR-421-3p, mmu-miR-497a-5p and mmu-miR-532-3pon the sequences of the linc/circRNAs (Fig EV2). LncRNAs in the cytoplasm may function as miRNA sponge. Thus, we analyzed the subcellular localization of the linc/circRNAs. Gapdh and Xist were checked as control. The results showed that the linc/circRNAs were mainly located in the cytoplasm (Fig 5D). Then, we wonder whether knockdown of the linc/circRNAs could lead to more the Oct4-miRNAs interacting with Oct4 mRNA, and performed RIP to captured Oct4 mRNA using the biotin-labeled miRNAs after the linc/circRNAs knockdown. We observed each of the linc/circRNA knockdown resulted in more enrichment of Oct4 mRNA comparing to control (Fig 5E). These results show that the linc/circRNAs function as Oct4-miRNAs sponges to sustain high expression of exogenous Oct4 during reprogramming.

**Figure 5.**
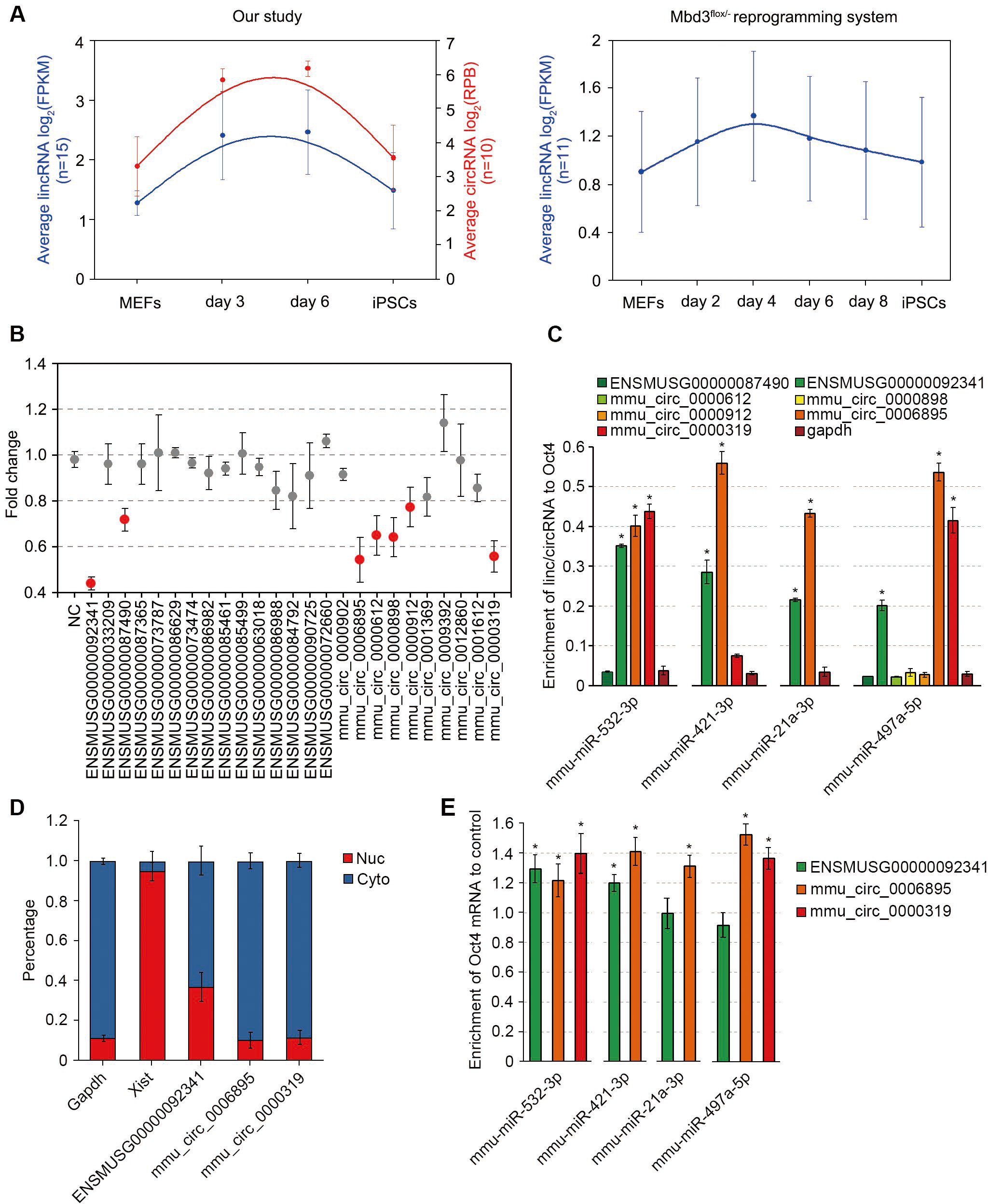
Linc/circRNAs sponge miRNAs targeting Oct4. A Expression patterns of linc/circRNAs which have complementary binding sites with the four Oct4-miRNAs. B Fold change of Oct4 mRNA with knockdown of the linc/circRNAs as indicated. Down-regulation of the linc/circRNAs highlighted in red significantly suppressed Oct4 expression by at least 20% (*p* <0.001). C RIP assay for detecting the interaction of the Oct4-miRNAs with the specific linc/circRNAs. Biotin-labeled miRNAs were transfacted into MEFs at a final concentration of 20 nM, and RIP assay was performed using day 3rMEFs. The enrichment of lincRNAs or circRNAs in bound fractions was evaluated by qRT-PCR analysis normalized to Oct4 mRNA. Gapdh was analyzed as negative control. D Subcellular localization analysis of the linc/circRNAs. Nuc, nucleoplasm; Cyto, cytoplasm. Gapdh and Xist act as cyto and nuc control, respectively. E Effect of the linc/circRNAs knockdown on interaction of the Oct4-miRNAs with Oct4 mRNA. RIP assay was performed using biotin-labeled miRNAs after knockdown of each of the linc/circRNAs as indicated. Scrambled siRNA non-targeting the linc/circRNAs was used as control. The error bars represent s.e.m.* indicates p <0.001.

### The linc/circRNAs can improve the reprogramming efficiency and the quality of iPSCs

The efficiency to derive AP positive clones was about 1.31% using the secondary reprogramming system (Fig 6A), and the iPSCs could give rise to chimeric pups and contributed to germline transmission (Table 1). Using the reprogramming system, we investigated whether the linc/circRNAs have any functional effects on reprogramming. To this end, we transfected each of the LNA-siRNAs targeting the linc/circRNAs in MEFs respectively, and we did not find any significant changes about the formation of alkaline phosphatase (AP)-positive colonies at day 17 post-OSKM induction. However, when the mixture of the LNA-siRNAs (si-mix) were transfected, we found that the colony formation was significantly decreased (Fig 6A) and only two cell lines were obtained. When additional Oct4 was added back (si-mix+Oct4-addback) by infecting MEFs using lentiviral Oct4, the number of AP-positive colonies was greatly enhanced (Fig 6A), indicating the low reprogramming efficiency is related to the decreased expression of exogenous Oct4induced by knockdown of the linc/circRNAs. Low level of exogenous Oct4 leads to generation of iPSCs with aberrant DNA methylation of the Dlk1-Dio3 locus and low capacity in chimeric mice [33]. Thus, we checked the methylation status of the Dlk1-Dio3 locus. Control iPSCs showed normal, 50%-60% DNA methylation, whereas the linc/circRNAs-knockdowned cells, especially the si-mix cells, showed DNA hypermethylation, and Oct4-addback could rescue the methylation defect (Fig 6B). Consistent with the hypermethylation pattern, all the miRNAs in the locus showed significantly lower expression in the linc/circRNAs-knockdowned cells comparing to the control iPSCs. Also, the expressions could be rescued by Oct4-addback (Fig 6C). In addition, we tested the *in vivo*-developmental potency of the iPSCs by injection into diploid blastocysts. In contrast, only one si-mix cell line could produce chimeric mice with the low extent of chimaerism and fail to support germline transmission (Table 1),whereas, all three Oct4-addback cell lines produced chimeric pups and one cell line could contribute to germline transmission (Fig 6D; Table 1). The results indicate that the linc/circRNAs can improve the reprogramming efficiency and the quality of iPSCs by sponging miRNAs targeting Oct4.

**Table 1.** *In vivo*-developmental potency of iPSCs

| **Cell lines** | **Blastocysts**<br>**injected** | **Chimeric/Total**<br>**pups** | **Chimaerism**<br>**(%)** | **Germline**<br>**transmission** |
| --- | --- | --- | --- | --- |
| **Con. Line 1** | 44 | 9/16 | 20-80 | yes |
| **Con. Line 2** | 48 | 7/22 | 40-70 | yes |
| **si-mix line 1** | 60 | 2/34 | 10, 20 | no |
| **si-mix line 2** | 62 | 0/26 | 0 | no |
| **si-mix+Oct4 line 1** | 46 | 8/22 | 30-80 | yes |
| **si-mix+Oct4 line 2** | 60 | 9/31 | 10-60 | no |
| **si-mix+Oct4 line 3** | 68 | 8/26 | 20-50 | no |

Note: The extent of chimaerism was estimated on the basis of coat color.

**Figure 6.**
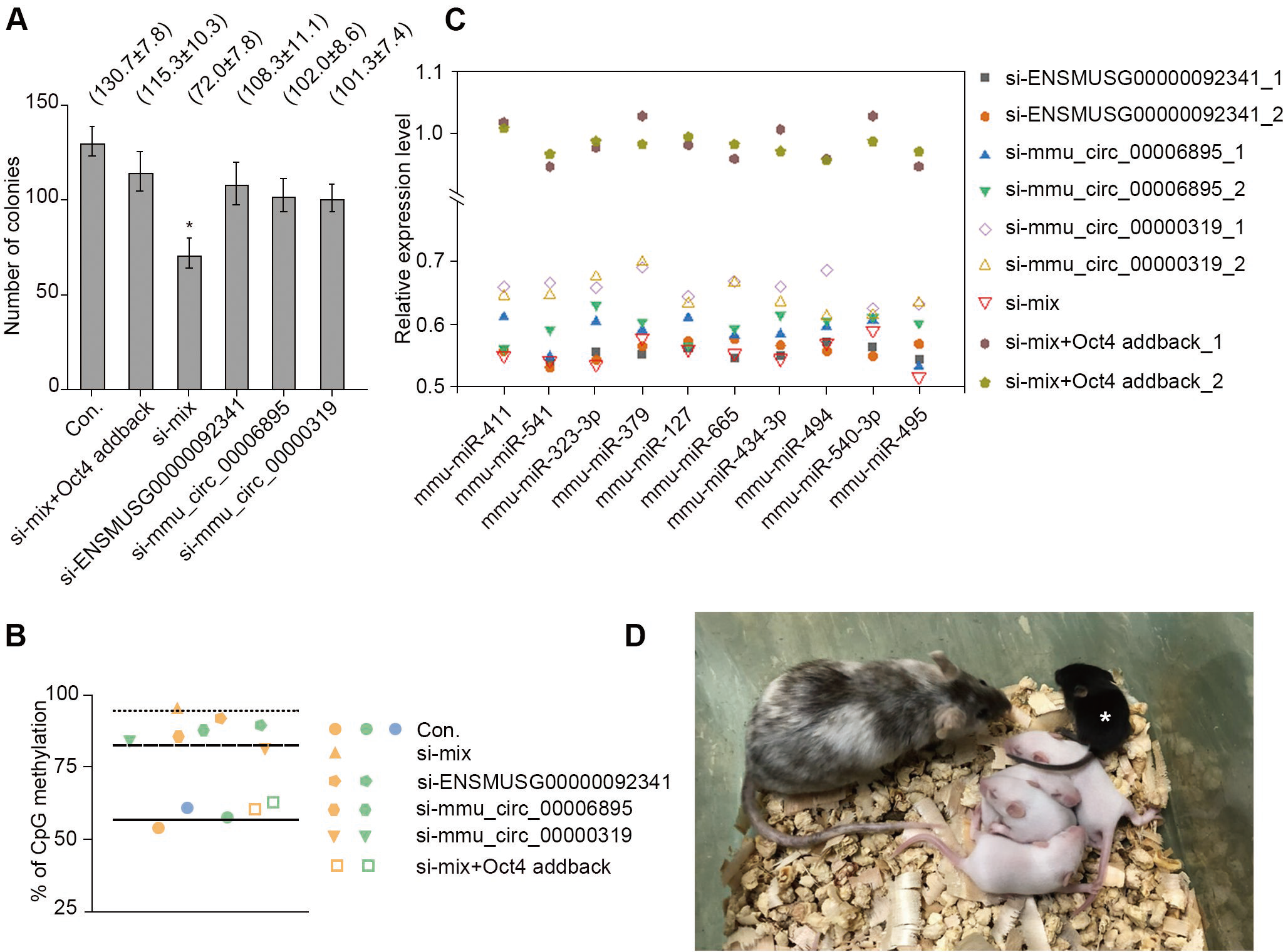
The linc/circRNAs can improve iPSCs reprogramming. A Effect of knockdown of the linc/circRNAs on formation of AP-positive iPSC colonies. MEFs were obtained from E13.5 embryos carrying a single copy of dox-inducible expression of the four transcription factors. AP staining was performed at day 17 after doxycycline addition. Before dox induction, 10 000 cells were placed in each well in a 6-well plate. Data represented as mean ± SD, *n* = 3. \**p* <0.001. B DNA methylation analysis on the Dlk1-Dio3 locus in cell lines. C Expression of the Dlk1-Dio3 locus encoded miRNAs in cell lines. Expression levels were normalized to control iPSCs. Con.: Cells transfected with negative control siRNA; si-mix: Cells transfected with mix of siRNAs targeting the linc/circRNAs; si-mix+Oct4 addback: Cells transfected with mixture of siRNAs targeting the linc/circRNAs and infecting by retroviral Oct4. Each shape represents a cell line. D Images of chimeras obtained from si-mix+Oct4 addback iPSCs and its F1 progeny resulting from germline transmission. *pup derived from the germline-competent iPSCs.

## Discussion

Herein we investigated the prevalence and properties of lncRNAs as ceRNA during reprogramming by experimental and bioinformatic approaches. We focused on the analysis of linc/circRNAs with the proposed role in the regulation of cell states from differentiation to pluripotency. Our data reveal the linc/circRNAs can sponge the miRNAs that target Oct4 mRNA and regulate expression of exogenous Oct4, and further improve reprogramming efficiency and quality of iPSCs (Fig 7). In the context of this critical cellular transition when repertoires of miRNAs are limited, the impact of linc/circRNAs on transcriptional programs is likely to be greater than in differentiated cells. Environmental or cellular stress, for example upon starvation or infection, may also offer similar opportunities for strong effects of ceRNA.

**Figure 7.**
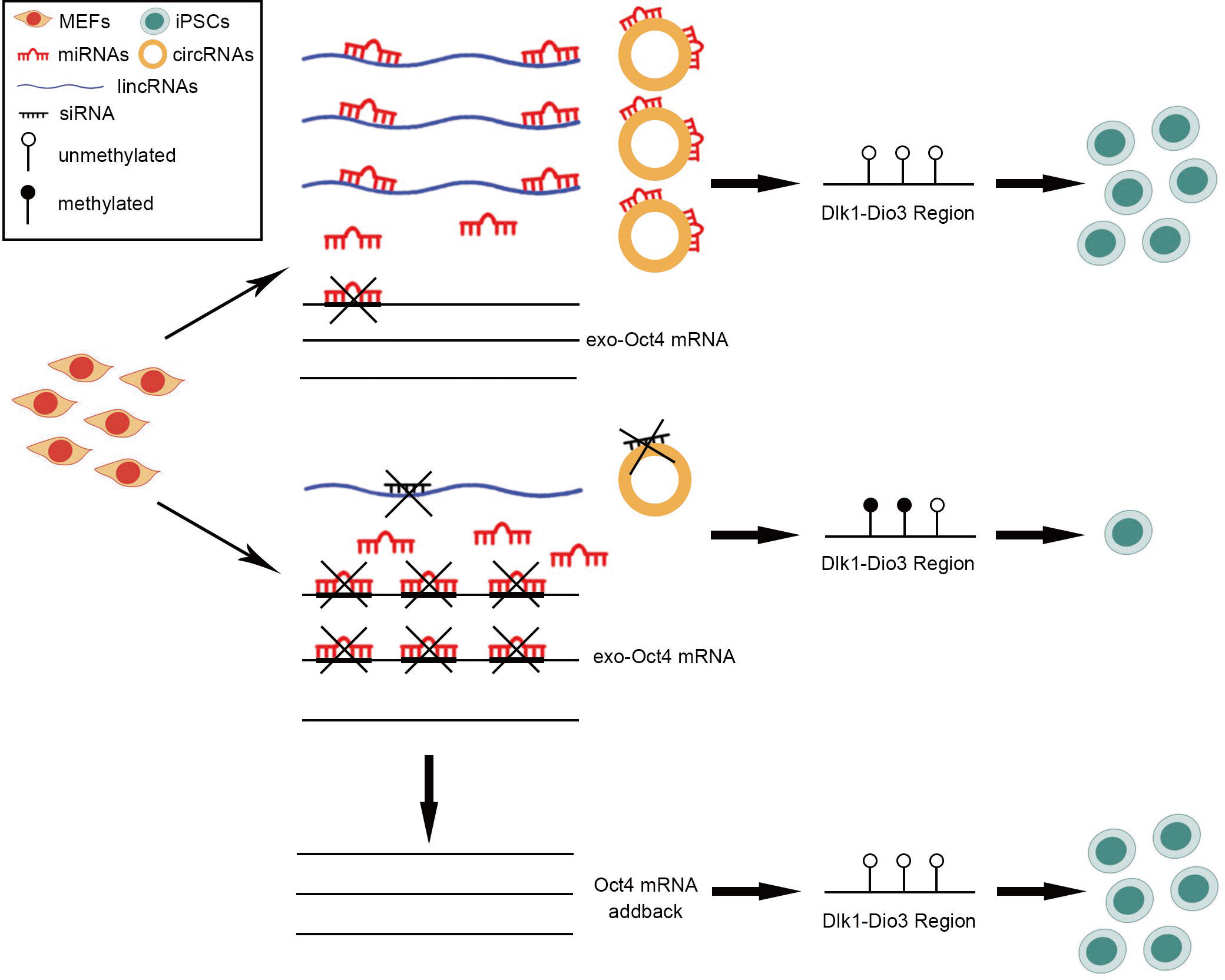
Model depicting linc/circRNAs as miRNAs sponges sustain high expression of exogenous Oct4 during reprogramming. During reprogramming, the miRNAs targeting on Oct4 mRNA from MEFs are highly expressed, however, the expression of exogenous Oct4 is not suppressed. Linc/circRNAs which have complementary binding sites with the miRNAs targeting Oct4 are also highly expressed and can counteract the activities of the miRNAs as sponges. The knockdown of the linc/circRNAs results in down-regulation of Oct4 expression, the imprinting defect at the Dlk1-Dio3 locus, decreasion of reprogramming efficiency, and low-grade chimera forming iPSCs, and those can be rescued by Oct4-addback. In summary, the linc/circRNAs can improve reprogramming efficiency and quality of iPSCs by sponging miRNAs targeting Oct4 during reprogramming.

We integrated the data of ribo-minus RNA-seq and small RNA-seq from cells collected during the course of reprogramming and fully reprogrammed iPSCs. During reprogramming, OSKM induction causes drastic changes of transcriptome [23, 27, 34]. It leads to inactivation of genes responsible for the identity of the parental cells and the activation of genes that are crucial for the establishment of the lineage of interest [23]. Similarly, in our study, we observed initial silencing of fibroblast genes and their transcriptomic activity was gradually replaced with switching-on of genes associated with pluripotency. However, we found the pool of miRNAs from MEFs was not changed remarkably during the course of reprogramming, indicating the reprogramming of miRNA expression pattern is a latter event. To ensure the successful reprogramming process, the activity of miRNAs targeting genes that need to be activated or up-regulated must be reduced. Indeed, we identified many linc/circRNAs, which share miRNA binding sites with mRNAs of pluripotent transcription factors, were highly expressed, suggesting that ceRNAs could be beneficial for reprogramming via miRNA-mediated mechanism.

LncRNAs proposed to act as sponges for miRNAs to modulate gene expression [4, 35]. In mouse ESCs, lots of lncRNAs have been predicted to compete for miRNAs with mRNAs [4]. However, it has not yet been experimentally validated as ceRNAs, especially circRNAs. Previous study has shown that linc-ROR regulates Oct4, Nanog, and Sox2 to sustain self-renewal of human ESCs by sponging miRNAs [11]. In the study, we identified that linc/circRNAs could serve as ceRNAs during reprogramming. We evidenced one lincRNA and two circRNAs could sustain high expression of core transcription factor Oct4 by sponging miRNAs during reprogramming, and they ultimately improved the reprogramming efficiency and the quality of iPSCs. During reprogramming, the activation of endogenous pluripotent genes happens at late stage [27, 36]. So, here, the ceRNA mechanism is involved in the regulation of exogenous Oct4 expression. Whether this mechanism regulates the endogenous pluripotent gene expression during reprogramming remains to be further determined. Together with previous studies [11], we suggest that the ceRNA network plays an important role on establishment and maintenance of pluripotency during reprogramming and in pluripotent stem cells.

In summary, our results point out a high prevalence of miRNA-mediated interactions between their mRNA targets and lncRNAs, proposing that this mechanism of ceRNAs indeed functions on regulation of gene expression, particularly in the context of cell-fate transitions.

## Material and mathods

### Animals

Mouse were housed and prepared according to the protocol approved bythe Institutional Animal Care and Use Committee of Northeast Agriculture University (Protocol number: IACUC-02-005) and the IACUC of Weill Cornell Medical College (Protocol number: 2014-0061).

### iPSCs reprogramming and high throughput RNA-seq

For the secondary reprogramming system, MEFs were obtained from E13.5 embryos from C57BL/6 mouse carrying a single copy of dox-inducible expression of OSKM. The MEFs within two passages were split when they reached 80–90% confluence, placed at 5000 cells/cm^2^. Before dox induction, 10 000 cells were placed in each well in a 6-well plate. 2ug/ml doxycycline was added to induce over-expression of OSKM. Reprogramming was performed in KSR medium (DMEM/F12 (Gibco) supplemented with 15% Knockout Serum Replacement (Gibco), 2 mM L-glutamine (Gibco), 1 × Pen/Strep (Gibco) 100 uM MEM non-essential amino acids (Gibco), 100 uM b-mercaptoethanol (Gibco), and 1000U/mL LIF (ESGRO, Millipore), 3 uM CHIR99021 (Stemgent) and 0.5× N-2Supplement (Gibco)). AP staining was performed at day 17 after doxycycline addition. Total RNA was isolated from the cells at day 0, day 3 and day 6 post-OSKM induction and the fully reprogrammed iPSCs using Trizol RNA extraction reagent (Thermo Fisher), following the manufacturer’s protocol. The concentration and purity of total RNA were assessed on Nanodrop2000 (Thermo Fisher Scientific, MA, USA), and the integrity of total RNA was evaluated with 2% agarose gel electrophoresis and RNA Nano 6000 Assay Kit of the Bioanalyzer 2100 system (Agilent Technologies, CA, USA). The cDNA libraries of long chain RNAs (mRNA/lncRNA/circRNA) and small RNAs were generated from 10 μg and 2 μg of total RNA, respectively. rRNA was removed using Epicentre Ribo-zero rRNA Kit (Epicentre, USA). Sequencing libraries were generated using NEBNext^®^ Ultra™ RNA Library Prep Kit for Illumina^®^ (#E7530L, NEB, USA) and NEBNext^®^ Multiplex Small RNA Library Prep Set for Illumina^®^ (#E7300L, NEB, USA), respectively. Sequencing was performed using Illumina HiSeq^TM^ 2000 platform by the Genomics Facility at Cornell Biotechnology Resource Center.

### Quantification of miRNA abundance and prediction of miRNA response elements

Clean sequencing reads were obtained by removing from the raw data reads containing more than one low quality (Q-value ≤ 20) base, with 5’ primer contaminants, without 3’ primer, without the insert, with poly A and shorter than 18nt. Clean reads were aligned with the reference genome (ftp://ftp.ensembl.org/pub/release-92/fasta/mus_musculus/dna/Mus_musculus.GRCm38.dna.toplevel.fa.gz), the Rfam database 13.0 [37] and the RepBase [38] to identify and remove rRNA, scRNA, snoRNA, snRNA, tRNA, repeat sequences, and fragments from mRNA degradation. The remaining reads were searched against miRBase 22.0 [39]to identify known miRNAs in mouse. The possible MREs of known miRNAs were predicted by RNA22 [32], an miRNA target prediction tool, with default parameter setting and the binding energy cutoff was less than or equal to −20 KCal/Mol. The numbers of MREs per kb length of each up-regulated or down-regulated genes and the average numbers of MREs in each groups were calculated. The miRNA expression level was calculated and normalized using RPM (reads per million). We identified significant differentially expressed miRNAs with a false discovery rate (FDR) < 0.05 and fold change ≥2 threshold by DEGseq [40].

### Identification of linc/circRNAs and expression analysis

Pre-processing of raw sequencing data were performed using FASTX-Toolkit with default parameters by removing low quality reads (More than 20% of the bases qualities are lower than 10), reads with adaptors and reads with unknown bases (N bases more than 5%). Clean reads were mapped to the mouse reference genome (ftp://ftp.ensembl.org/pub/release-92/fasta/mus_musculus/dna/Mus_musculus.GRCm38.dna.toplevel.fa.gz). Unmapped reads were collected and the software CIRI [41] and find_circ [17] were used to identify circRNAs with default options, and circBase IDs was used to indicate known circRNAs. Counts of identified circRNA reads were normalized by RPB (junction reads per billion mapped reads) method. For lincRNAs, the reconstruction and identification of transcripts were performed by mapped reads with StringTie [42] and cuffcompare [43]. Known lincRNAs were acquired by gene annotations for comparing the assembled transcripts with NCBI, Ensembl and UCSC mouse known genes. The expression level of lincRNA was calculated using FPKM (Fragments Per Kilobase of transcript per Million mapped reads) with the software RSEM [44]. Transcripts with a false discovery rate (FDR) <0.05 and fold change ≥2 were then identified as significant differentially expressed lincRNA or circRNA using DEGSeq [40].

### Experimental validation of miRNA activity and interaction between mRNAs and linc/circRNAs

The miRNAs mimics and inhibitors, and linc/circRNAs siRNAs were synthesized by Exiqon, and were transfected in triplicates into MEFs using Lipofectamine RNAi MAX Reagent (Invitrogen) according to the manufacturer’s guidelines. rMEFs were collected at day 3 post-OSKM induction, RNAs were extracted and checked by qRT-PCR for gene expressions. RIP was performed using Magnetic RNA-Protein Pull-Down Kit (Pierce, #20164) according to the manufacturer’s protocol. Briefly, the 3’end biotinylated miRNAs mimics were transfected into MEFs at a final concentration of 20 nM for 1 day. The biotin-coupled RNA complex was pull-downed by incubating the cell lysates with streptavidin-coated magnetic beads. The abundance of linc/circRNAs in bound fractions was evaluated by qRT-PCR analysis normalized to Oct4 mRNA. Gapdh was analyzed as negative control. The TetO-FUW-Oct4 plasmid (Add gene plasmid 20323) was used for Oct4-addback, and 10 000 cells infected with a dose of 5×10^8^ cfu of virus for 3 h.

### Methylation analysis of the Dlk1-Dio3 locus

Genomic DNA isolation from cells and bisulfite pyrosequencing were performed as follows. Briefly, genomic DNA of iPSCs was extracted with the Gentra Pure gene Kit (Qiagen) according to the manufacturer’s instructions and quantified using a Nano Drop 2000 spectrophotometer. Bisulfite conversion of 500 ng genomic DNA per sample was performed with the EpiTect Bisulfite Kit (Qiagen) according to the manufacturer’s specifications. Quantification of DNA methylation was carried out by PCR of bisulfite-converted DNA and pyrosequencing. PCR and sequencing primers for bisulfite pyrosequencing were designed using the Pyrosequencing Assay Design Software 2.0 (Qiagen). For pyrosequencing, a PyroMark Q96 ID instrument (Qiagen) and PyroMark Gold Q96 reagents (Qiagen) were used. Data were analyzed using the PyroMark CpG Software 1.0.11 (Qiagen).

### Statistical analysis

All data are presented as mean ± s.e.m. Differences between groups were tested for statistical significance using Student’s *t-*test or χ^2^-test. Statistical significance was set at *p <0.01*.

## Acknowledgements

This work was supported by the Natural Science Foundation of Heilongjiang Province (C2018032) (QK), the Starr Foundation Tri-Institutional Stem Cell Core and Derivation grant (DW), and The National Key Research and Development program of China-Stem cell and Translational Research (2016YFA0100200) (ZL).

## Author Contributions

QK and DW conceived and designed the experiments; QK and JZ conducted the experiments and collected data; XZ, KZ, HZ and XP analyzed RNA-seq and small RNA-seq data; QK and XZ wrote the manuscript; DW and QK discussed and edited the manuscript; ZL and DW provided resource and reviewed the manuscript.

## Data availability

RNA-seq and small RNA-seq data: Gene Expression Omnibus GSE116684 (https://www.ncbi.nlm.nih.gov/geo/query/acc.cgi?acc=GSE116684)

